# Queuine salvaging in the human parasite *Entamoeba histolytica*

**DOI:** 10.1101/2022.06.21.496972

**Authors:** Lotem Sarid, Jingjing Sun, Jurairat Chittrakanwong, Meirav Trebicz-Geffen, Peter C. Dedon, Serge Ankri

**Affiliations:** Department of Molecular Microbiology, Ruth and Bruce Rappaport Faculty of Medicine, Technion, Haifa, Israel; Department of Biological Engineering, Massachusetts Institute of Technology, Cambridge, Massachusetts, USA; Laboratory of Biotechnology, Chulabhorn Research Institute, 54 Kamphaeng Phet 6 Road, Talat Bang Khen, Lak Si, Bangkok 10210, Thailand

## Abstract

Queuosine (Q) is a naturally occurring modified nucleoside that occurs in the first position of transfer RNA anticodons such as Asp, Asn, His, and Tyr. As eukaryotes lack pathways to synthesize queuine, the Q nucleobase, they must obtain it from their diet or gut microbiota. Previously, we described the effects of queuine on the physiology of the eukaryotic parasite *Entamoeba histolytica* and characterized the enzyme EhTGT responsible for queuine incorporation into tRNA. At present, it is unknown how *E. histolytica* salvages Q from gut bacteria. We used liquid chromatography–mass spectrometry (LC–MS) and N-acryloyl-3-aminophenylboronic acid (APB) PAGE analysis to demonstrate that *E. histolytica* trophozoites can salvage queuine from Q or *E. coli* K12 but not from the modified *E. coli* QueC strain, which cannot produce queuine. We then examined the role of EhDUF2419, a protein with homology to DNA glycosylase, as queuine salvage enzyme in *E. histolytica*. We found that glutathione S-transferase (GST)-EhDUF2419 catalyzed the conversion of Q into queuine. Trophozoites silenced for EhDUF2419 expression are impaired in their ability to form Q-tRNA from Q or from *E. coli*. We also observed that Q partially protects control trophozoites from oxidative stress (OS), but not siEhDUF2419 trophozoites. Overall, our data reveal that EhDUF2419 is central for the salvaging of queuine from bacteria and for the resistance of the parasite to OS.

## Introduction

In many parts of the world, poor sanitation and unsafe hygiene practices are causing amebiasis to spread. The World Health Organization estimates that 50 million people in India, Southeast Asia, Africa, and Latin America suffer from amebic dysentery and amebiasis each year, resulting in at least 100,000 deaths. Amebiasis is primarily transmitted through ingesting contaminated food or water containing *E. histolytica* cysts. After the cyst form has been swallowed by the host, excystation occurs in the intestinal lumen, followed by colonization of the large intestine by the trophozoites. These next divide and encyst; both trophozoites and cysts are excreted in stools. *E. histolytica* trophozoites reside in the colon as a non-pathogenic commensal in most infected individuals (90% of infected individuals are asymptomatic). For unknown reasons, the trophozoites can become virulent and invasive, cause amebic dysentery, and migrate to the liver via the portal veins, where they cause hepatocellular damage. No vaccine against amebiasis currently exists; the drug of choice for treating amebiasis is metronidazole, which, however, may have severe side effects. Additionally, some clinical strains of *E. histolytica* are less sensitive to metronidazole, suggesting the emergence of metronidazole-resistant strains [1]. RNA modifications are emerging as an essential means to maintain the cell life cycle in numerous contexts, ranging from infection to neuropathologies and cancer. More than 100 RNA chemical modifications are known to date, addressing all RNA species. Nevertheless, RNA modifying enzymes have not yet been exploited as drug targets.

Queuosine (Q) and its glycosylated derivatives occur in position 34 of the anticodon of tRNA with G_34_U_35_N_36_ in the anti-codon loop of eubacteria and eukaryotes except for *Saccharomyces cerevisiae* [2,3]. Q is highly conserved and found in plants, fishes, insects and mammals. While many bacteria can synthesize queuine (the nucleobase of Q) *de novo*, salvage of the prokaryotic Q precursors preQ_0_ and preQ_1_ has recently be reported [4]. Eukaryotes are not capable of Q synthesis and rely on salvage of the queuine base as a Q precursor either by nutrition or by the intestinal bacterial flora [5-7]. The tRNA-guanine transglycosylase (TGT) is the main enzyme responsible for the formation of Q into the wobble position of the anti-codon loop. The enzyme exchanges G34 for the precursors. The cyclopentendiol moiety is synthesized at the level of tRNA from unknown precursors and enzymes in both eubacterial and eukaryotic species. The crystal structure of hTGT in its heterodimeric form and in complex with a 25-mer stem loop RNA has been recently established [8]. The detailed analysis of its dimer interface and interaction with a minimal substrate RNA indicates that one base only, guanine 34 or queuine, can simultaneously reside at the active site in support to a “ping-pong” mechanism that has already been proposed for *E. coli* TGT [9]. Regarding hQTRTD1, the authors proposed that it could serve to anchor the TGT enzyme in the compartmentalized eukaryotic cell [8]. Based on the annotation of the *E. histolytica* genome, a homolog of *h*QTRT1 and *h*QTRTD1 exists in *E. histolytica*, namely *Eh*QTRT1 (XP_656142.1) and *Eh*QTRTD1 (XP_652881.1). Our previous work has significantly contributed to our understanding of *E. histolytica* tRNA-guanine transglycosylase (TGT) (EhTGT) and of the regulation of *E. histolytica’*s virulence by queuine [10]. We found that EhTGT is a heterodimer composed of *Eh*QTRT1 and *Eh*QTRTD1. EhTGT is catalytically active and it incorporates queuine into *E. histolytica* tRNAs. The presence of Q in tRNA^Asp^_GUC_ stimulates its methylation by Ehmeth, a Dnmt2-type multisubstrate tRNA methyltransferase, at the C38 position. Queuine does not affect the growth of the parasite, it protects the parasite against oxidative stress (OS) and it antagonizes the negative effect that OS has on translation by inducing the expression of genes involved in OS response, such as heat shock protein 70 (Hsp 70), antioxidant enzymes, and enzymes involved in DNA repair. On the other hand, queuine impairs *E. histolytica* virulence determined in mouse model of amebic colitis by downregulating the expression of genes previously associated with virulence, including cysteine proteases, cytoskeletal proteins, and small GTPases. Silencing of EhQTRT1 expression prevents the incorporation of queuine into tRNAs, impairs the methylation of C38 in tRNA^Asp^_GUC_, inhibits the growth of the parasite, impairs its resistance to OS and its cytopathic activity.

Information about how Q is salvage by eukaryotic organisms is scanty. In mammalian cells, queuine is generated from Q-5’-phosphate which suggests that salvage Q base is coming from tRNA degraded during the normal turnover process [11]. In the green algae *Chlorella pyrenoidosa* and *Chlamydomonas reinhardtii* an enzymatic activity that catalyzes the cleavage of Q has been identified but its nature is not known [12]. A search of genes that co-distribute with eukaryotic QTRT1 and QTRTD1 identified a potential Q salvage protein, DUF2419 [13]. The structural similarity of DUF2419 with DNA glycosylases suggests a ribonucleoside hydrolase activity. Indeed, genetic evidences support the role of DUF2419 as a Q salvaging enzyme in *Schizosaccharomyces pombe*, human, maize, and *Streptococcus thermophilus* [13]. Here, we used a genetic approach coupled LC-MS and APB PAGE analysis to demonstrate the EhDUF2419 is the enzyme that salvage Q in *E. histolytica*.

## Materials and Methods

### *E. histolytica* culture

*E. histolytica* trophozoites, the HM-1:IMSS strain (a kind gift of Prof. Samudrala Gourinath, Jawaharlal Nehru University, New Delhi, India), were grown and harvested according to a previously reported protocol [14]. The trophozoites were cultivated with Q (0.1 μM) (a gift from Prof. Peter C. Dedon, MIT, USA) or queuine (0.1 μM) (a gift from Prof. Klaus Reuter, University of Marburg, Germany) for three days.

### Transfection of *E. histolytica* trophozoites

The transfection of *E. histolytica* trophozoites was performed using a previously described protocol [15].

### Resistance of *E. histolytica* trophozoites to OS

Resistance to OS of trophozoites was determined by the eosin dye exclusion method [16].

### Growth rate of *E. histolytica* trophozoites

4x 10^4^ *E. histolytica* trophozoites were grown in 15-mL tube in TYI-S-33 medium at 37°C. The number of viable trophozoites were counted according to previously described protocol [16] after 24 and 48 hours of culture.

### Primer used in this study

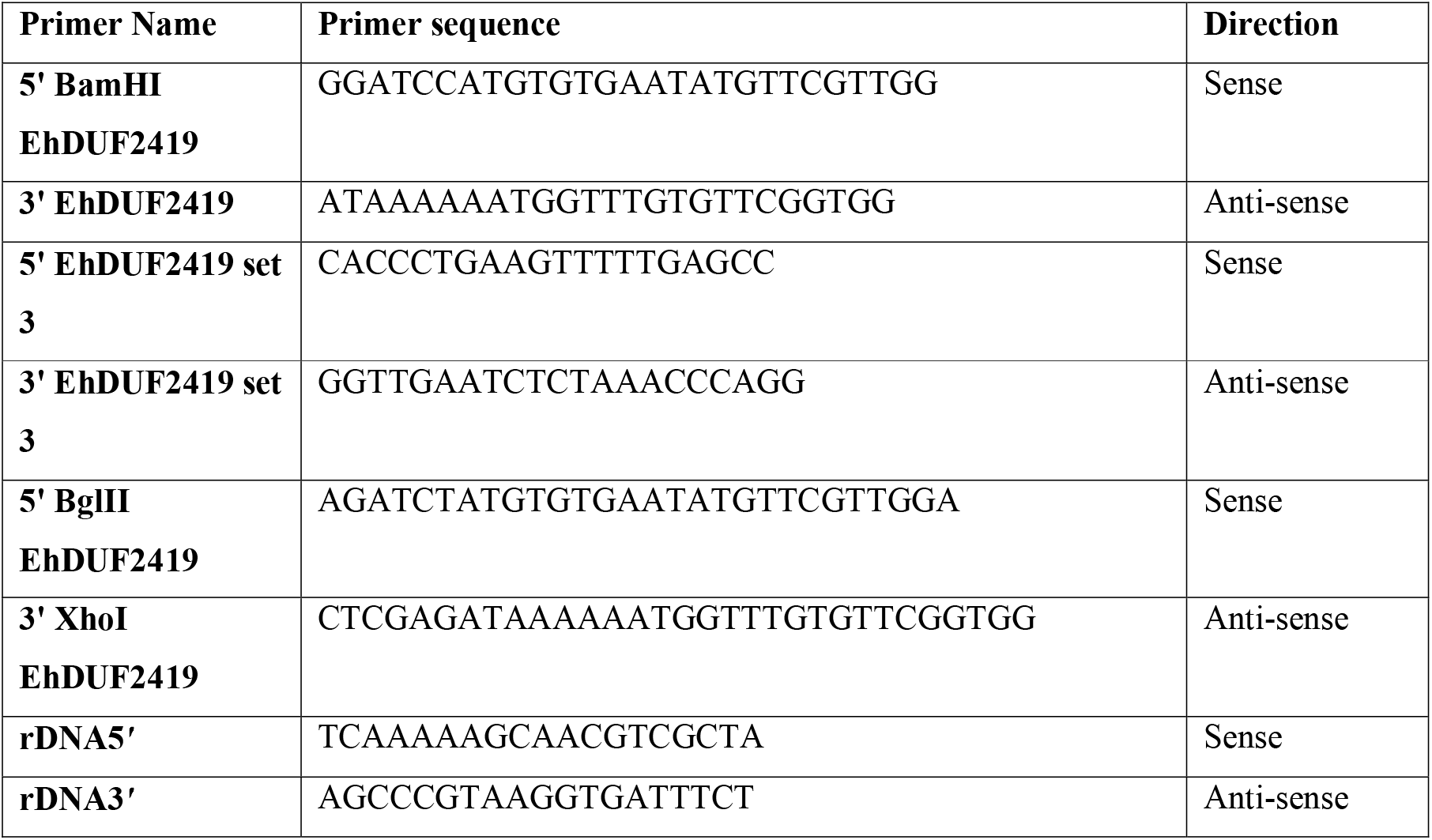

### Construction of GST-tagged EhDUF2419 vector

For construction of the pGEX-EhDUF2419 vector, EhDUF2419 was amplified from *E. histolytica* genomic DNA with the primers 5’ BamHI EhDUF2419 and 3’ EhDUF2419. The PCR product was cloned in a pGEM-T easy vector (Promega), digested with BamHI and NotI, and then subcloned into a pGEX-4T-1 (Pharmacia Biotech) vector to generate pGEX-EhDUF2419 vector.

### Construction of silenced EhDUF2419 vector

For construction of the siEhDUF2419 vector, EhDUF2419 was amplified from *E. histolytica* genomic DNA with the primers 5’ BglII EhDUF2419 and 3’ XhoI EhDUF2419. The PCR product was cloned in a pGEM-T easy vector (Promega), digested with BglII and XhoI, and then subcloned into a pEhEx-04-trigger vector containing a 142-bp trigger region (EHI_048660) (a kind gift of Tomoyoshi Nozaki, University of Tokyo, Japan) to generate siEhDUF2419 vector.

### Preparation of recombinant GST-tagged EhDUF2419

Recombinant EhDUF2419 was expressed as GST-tagged protein at *E. coli* BL21(DE3)pLysS competent cells, which were transfected with pGEX vector derived plasmids. The overnight culture was supplemented with Amp (100μg/ml) and grown at 37°C until the OD_600_ reached ~0.4. Synthesis of the GST-tagged protein complex was initiated by adding IPTG to the culture at a final concentration of 0.1mM. After an overnight incubation in the presence of IPTG at 16°C, the bacteria were harvested and lysed with lysis buffer (100mM KCl, 1mM DTT, 1mM PMSF, 0.1μg/mL Lysozyme, 0.1μg/mL Leupeptine, PBS) and set in ice for 30min. The cells were sonicated on a Sonics Vibracell VCX750 Ultrasonic Cell Disrupter (Labotal) and BugBuster protein extraction reagent (1:100 ratio) (Novagen) were added to complete the lysis. GST-tagged proteins were purified under native conditions on gluthatione-agarose resin (Sigma). The recombinant proteins were then washed 3 times with Buffer A (100mM KCl, 1% Triton, 1mM PMSF, PBS) and then 3 times with Buffer B (100mM KCl, 1mM PMSF, PBS). Next, the proteins were eluted with glutathione elution buffer [Tris HCl 50mM pH-9.6, L-glutathione reduced (Sigma) 10mM, 150mM NaCl]. Eluted proteins were resolved on 12% SDS gel and the gel was stained with silver (Pierce Silver Stain).

### Preparation of recombinant GST

Recombinant GST was expressed in *E. coli* BL21(DE3)pLysS competent cells, which were transfected with pGEX-4T-1 vector. The proteins were prepared according to the protocol described above.

### Enzymatic activity of EhDUF2419

2.5μg of recombinant GST or GST-tagged EhDUF2419 was incubated with 409 ng of Q in 30-μl HEPES-reaction buffer (100mM HEPES pH-7.3, 20mM MgCl_2_, 5mM DTT) at 37°C overnight. Next, GST or GST-EhDUF2419 was pulldown from the reaction with glutathione-agarose resin. The level of Q vs queuine in the samples was determined by LC-MS/MS as described below.

### RNA extraction

i. RNA extraction using Trizol-Total RNA was extracted using the TRI reagent kit according to the manufacturer’s instructions (Sigma-Aldrich).
ii. RNA extraction using Monarch Total RNA Miniprep kit-total RNA was extracted from *E. histolytica* trophozoites that were incubated with *E. coli* K12/ΔQueC (a kind gift of Prof. Valérie de Crécy-Lagard, University of Florida, USA) using the Monarch Total RNA Miniprep Kit (NEW ENGLAND BioLabs) according to the manufacturer’s instruction.

### Quantification of tRNA modification by LC-MS/MS

The method for tRNA modification quantification was modified from an established LC-MS/MS method, which include tRNA purification, tRNA hydrolyzation and LC-MS/MS analysis [17].

#### tRNA purification using HPLC

RNA samples were re-suspended with 50 μL of RNase-free water. Twenty microliter of each sample was injected to purify by HPLC (Agilent 1200 series). Agilent SEC-3 300-Å HPLC column (Agilent Technologies, cat. no. 5190-2511) was used to purify tRNA from total RNA with a temperature-controlled column compartment at 40 °C with 100 mM ammonium acetate aqueous phase at a flow rate of 1 ml/min. The tRNA peak was collected by Agilent automatic fraction collector and transferred into a 1.5 mL tube. All samples were dried by SpeedVac vacuum concentrators and re-dissolved with 50 μL of RNase-free water. Concentration of all samples was measured using nanodrop. Representatives of samples were used to tRNA purity and integrity using Agilent RNA 6000 nano kit with Agilent 2100 bioanalyzer instrument.

#### Hydrolysis of tRNA to nucleosides

Four-hundred nanogram of each sample was hydrolyzed in 96-well plate in 30 μL reaction volumn. The reaction mixture included 2.5 mM MgSO_4_, 5 mM Tris (pH 8.0), 0.1 μg/mL coformycin, 0.1 mM deferoxamine, 0.1 mM butylated hydroxytoluene, 0.083 U/μL benzonase, 0.1 U/μL calf intestinal alkaline phosphatase, 0.003 U/μL phosphodiesterase I, and 50 nM internal standard [^15^N]_5_-deoxyadenosine. The reactions were incubated at 37 °C for 6 h and used for LC-MS/MS analysis without further purification.

#### LC-MS/MS quantification analysis

LC-MS/MS analysis parameters were developed by using purchased and synthetic standards, including the LC gradient and the retention time, m/z of the transmitted parent ion, m/z of the monitored product ion, fragmentor voltage, and collision energy of each modified nucleoside. In brief, samples were loaded on a Phenomenex Synergi Fusion-RP C18 column (100 × 2.0 mm, 2.5 μm), and tRNA modifications were eluted with a gradient starting with 100% phase A (5 mM ammonium acetate, pH 5.3), followed by 0-10% phase B (acetonitrile) 0-10 min; 10%-40% phase B, 10-14 min; 40%-80% phase B, 14-15 min; 80%-90% phase B, 15-15.1 min; 90% phase B, 15.1-18 min at 35 degree and a flow rate of 0.35 ml/min. The HPLC column was coupled to an Agilent 6490 Triple Quad mass spectrometer with an electrospray ionization source in positive mode with the following parameters: gas temperature, 200 degree; gas flow, 11 l/min; nebulizer, 20 psi; sheath gas temperature, 300 degree; sheath gas flow, 12 l/min; capillary voltage, 1800 V; Vcharging, 2000 V. MRM mode was used for detection of product ions derived from the precursor ions for all the tRNA modifications with instrument parameters including the collision energy (CE) optimized for maximal sensitivity for the modifications.

### APB northern blotting for *E. histolytica* tRNA^His^_GUG_

APB gels were prepared and run with a few modifications according to Igloi and Kössel [18]. Briefly, 14□μg of total RNA was deacetylated in 100□mM Tris-HCl [pH 9] for 30□min at 37°C. RNA was ethanol precipitated and resuspended in 10□μl DEPC-treated water. Samples were then denatured for 10□min at 70°C and run at 4°C on Tris-acetate EDTA (TAE) buffer, 8 M urea, 15% acrylamide, and 5□mg/ml aminophenylboronic acid (Sigma) on Bio-Rad mini gels. The gel was run at 4°C at 75 V for 7-8□hours until the bromophenol blue reached the bottom of the gel. The gels were then stained with ethidium bromide in 1× TAE buffer for 20 min and then visualized for equal loading of samples. The gels were destained with ultrapure water for 20 min, and samples were transferred to a Nylon Hybridization Transfer membrane (PerkinElmer) by electrotransfer using 0.5× TAE as the transfer buffer for 45 min at 130mA. The membrane was cross-linked by UV using 120 mJ (Stratagene UV linker) and hybridized twice for 15 min each in 5□ml hybridization buffer (20□mM sodium phosphate buffer [pH-7], 300□mM NaCl, 1% SDS), followed by the addition of 150□μg/ml heat-denatured salmon sperm DNA (ssDNA) to the hybridization buffer and blocking for 1□h at 60°C. The membrane was then incubated with 15□pmol\mL of biotinylated tRNA probes prepared against *E. histolytica* tRNA^His^_GUG_ and incubated at 60°C for 16-18□hours. The membrane was then washed for 10 min with 5□ml wash buffer (20□mM sodium phosphate buffer [pH 7], 300□mM NaCl, 2□mM EDTA, 0.5% SDS) at 60°C and then incubated in hybridization buffer once at RT for 10 min. The membrane was then incubated in streptavidin-horseradish peroxidase (HRP) conjugate in 5□ml hybridization buffer (1:5,000) for 1 hour followed by two washes for 10 min. The membranes were incubated with enhanced chemiluminescence reagent (Advansta), and exposed in FUSION FX7 (Vilber).

### Quantitative-Real Time PCR

Total RNA was extracted from control (WT) or siEhDUF2419 trophozoites using TRI reagent (Sigma) and the amount of total RNA was quantified using nanodrop spectrophotometer (ThermoFisher scientific). Reverse transcription was performed using the RevertAid First strand cDNA synthesis kit (ThermoFisher scientific), according to the manufacturer’s instructions. qRT-PCR was performed using the qPCRBio SyGreen Mix Hi-ROX (PCR biosystems) according to the manufacturer’s instructions and run on the Real Time PCR QuantStudio3 (ThermoFisher scientific) using the following conditions – Initial denaturation step at 95□ °C for 2 minutes, 40 cycles of denaturation at 95□ °C for 5 seconds, followed by hybridization at 50□°C for 30 seconds. The melting curve was performed according to the following conditions: 95°C for 15 seconds, then 60°C for 1 minute, and finally at 95°C for 15 seconds. The relative fold change was calculated using the 2 ^^(−ΔΔCt)^ method [19]. The QRT-PCR values were normalized to the level of rDNA [20]. PCR amplification controls were performed for each primer to verify the formation of a single PCR product.

### Western blotting

Western blotting of total protein extract of *E. histolytica* trophozoites (40μg) was performed according to previously described protocol [10]. Briefly, the proteins were resolved on 12% SDS gel and electrotransferred to nitrocellulose membrane (Whatman, Protran BA83). The blots were blocked with 5% skim milk and then probed with mouse polyclonal-EhTGT antibody (1:1000) [10] for 16 hours at 4°C. Next, the blots were washed, probed with secondary antibody (Jackson ImmunoResearch) at room temperature for one hour, and developed using enhanced chemiluminescence reagent (Advansta).

## Results and discussion

### Salvage of Q by *E. histolytica*

In order to determine whether the parasite can salvage Q, *E. histolytica* trophozoites were cultivated in the presence of Q and the Q-tRNA level was determined by LC–MS (Fig. 1). In agreement with our previous data [10], we found that queuine is uptaken by *E. histolytica* trophozoites and incorporated into tRNA (Fig 1). When trophozoites were grown with Q, the levels of Q-tRNA is significantly higher than the level in trophozoites grown without Q. These results indicate that a transporter(s) mediate the uptake of Q or queuine inside the parasite. Recently, a queuine transporter, ypdP, has been identified in the pathogenic bacteria *Chlamydia trachomatis* ypdP [4]. However, we did not find a *C.trachomatis* ypdP homologue in *E. histolytica* by using BlastP [21]. Mediated transport of nucleoside in *E. histolytica* has been previously demonstrated [22] and a number of transmembrane proteins identified [23]. Parasitic protists possess nucleoside and nucleobase transporters that belong to the equilibrative nucleoside transporter (ENT) family, which has eleven membrane-spanning domains and occurs in animals and plants alike [24]. Three members of the ENT family namely (EHI_110730, EHI_017040 and EHI_1421500) have been identified among *E. histolytica* transmembrane proteins [23] and their role in the transport of queuine or Q is currently studied.

**Figure 1:**
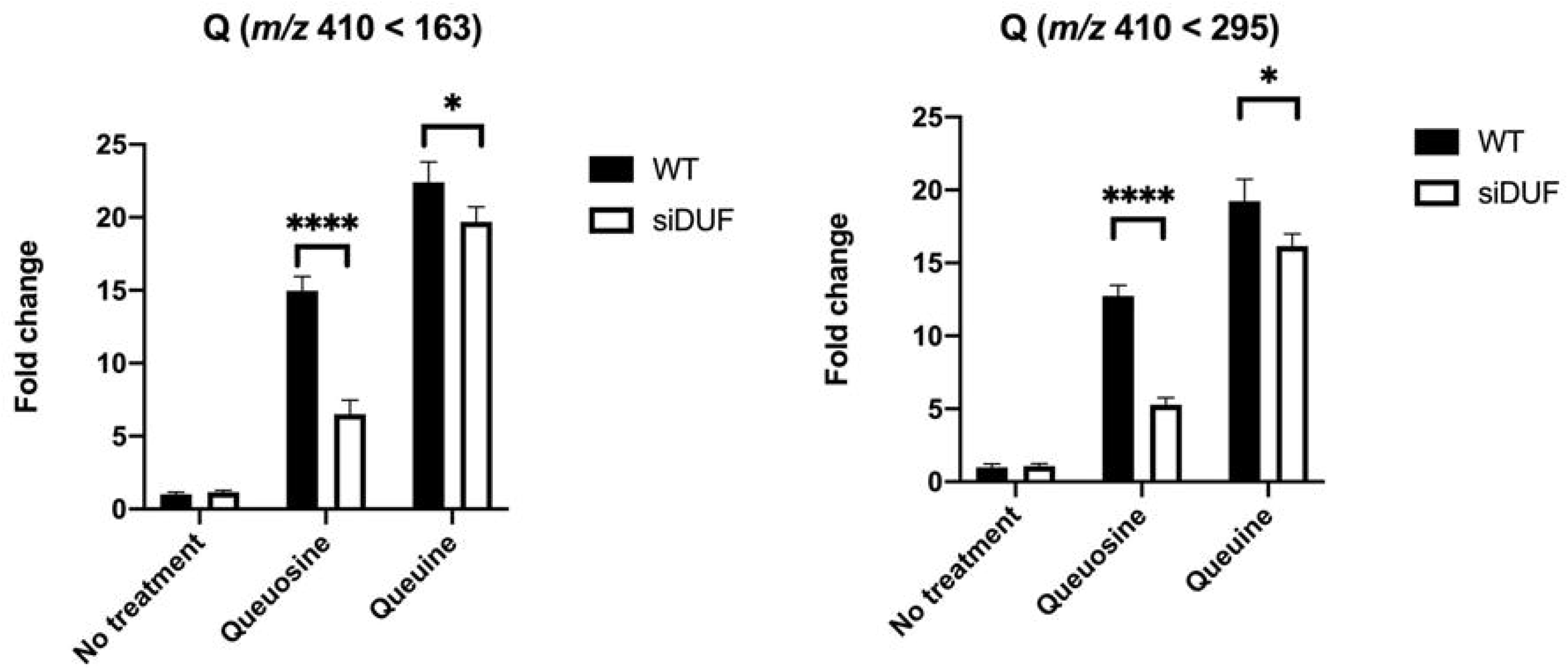
Q level change determined by LC-MS upon in trophozoites cultivated with queuosine or Q. The fold change is relative to wild type strain without any additive treatment. * indicates *P* value < 0.05, ****indicates *P* value < 0.0001.

In the large intestine, *E. histolytica* trophozoites feed on bacteria [25]. Consequently, we hypothesized that the parasite can salvage queuine directly from ingested bacteria. The hypothesis was tested by feeding *E. histolytica* trophozoites with *E. coli* K12 that served as a Q-donor bacteria. [26]. According to LC-MS data, trophozoites can salvage queuine from *E. coli* K12 (Fig 3). We confirmed that *E. coli* K12 is the source of queuine as *E. histolytica* trophozoites were unable to salvage queuine from *E. coli* ΔQueC, a mutant which is unable to synthesize queuine [27] (Fig 3). The conclusions drawn from LC-MS data are further supported by studying the level of Q-tRNA^His^_GUG_ by APB polyacrylamide gel analysis (Fig 2&4). The mammalian gut is crowded with microorganisms fighting for nutrients and for survival [28]. Competition for Q also occurs in the gut and recently, two Q salvage pathways have been characterized in pathogenic and commensal bacteria [4]. *E. histolytica* will be exposed to this Q competition inside the human gut. Data shown in Fig 3&4 indicate that the parasite has an advantage over its prokaryotic competitors because it phagocytoses bacteria, the main source of Q. Some bacteria like *Lactobacillus ruminus* are preferred as a nutritional source by *E. histolytica* over other gut bacteria [25]. There is a possibility that *L. ruminus* may offer the parasite an important source of Q. *L. ruminus* encodes a TGT enzyme in its genome (WP_014073827.1), supporting this hypothesis.

**Figure 2.**
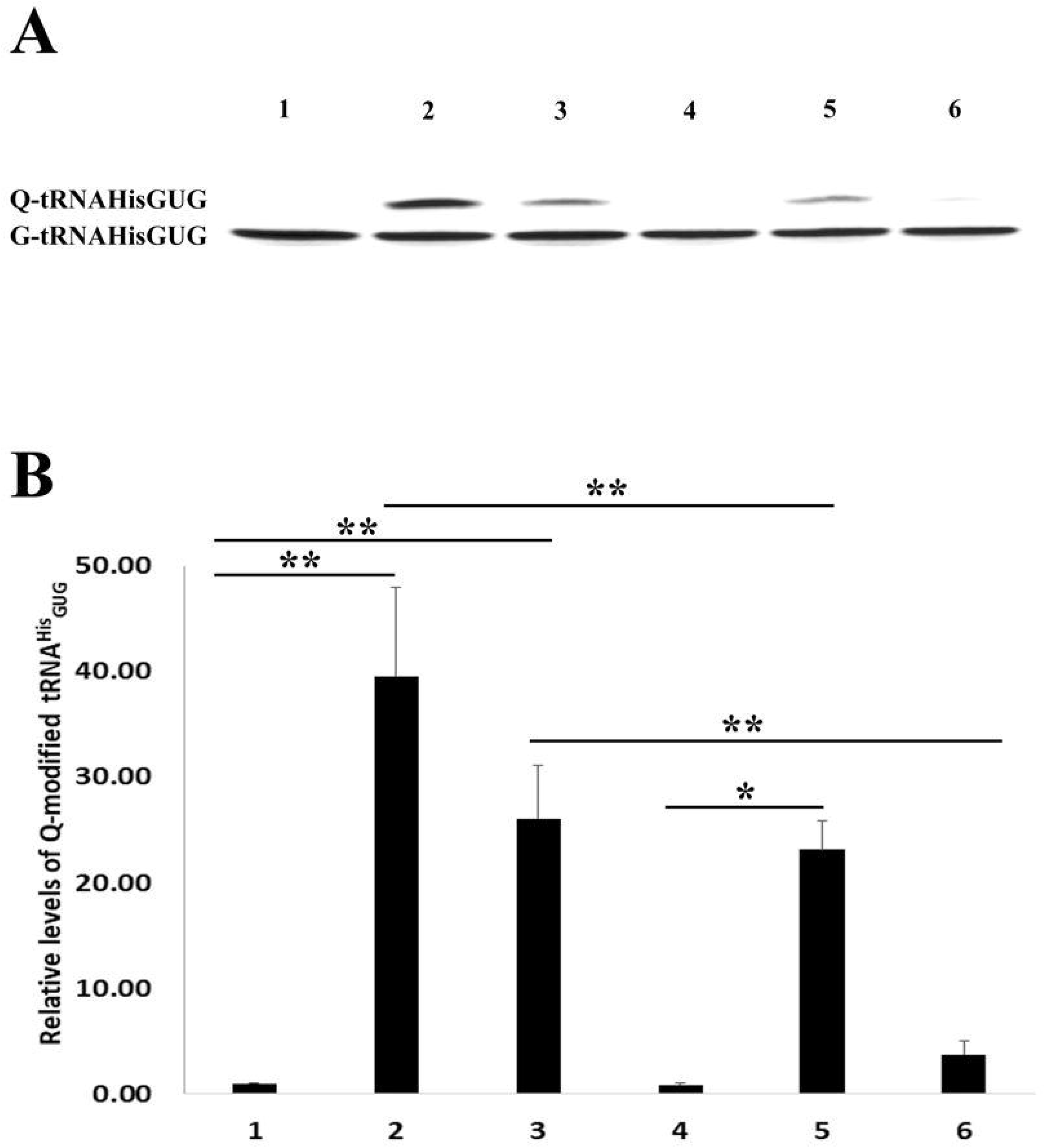
APB northern blot analysis of tRNA^His^_GUG_ in control (WT) and siEhDUF2419 trophozoites following cultivation with queuine or Q. Control (WT) or siEhDUF2419 trophozoites were cultivated in the presence of 0.1μM queuine or Q for 3 days. (1) Wild-Type trophozoites (2) queuine-treated WT trophozoites (3) Q-treated WT trophozoites (4) siEhDUF2419 trophozoites (5) queuine-treated siEhDUF2419 trophozoites (6) Q-treated siEhDUF2419 trophozoites. The data represent two independent experiment that were repeated twice. p value<0.05 by an unpaired Student *t* test (* ≤0.05, ** ≤0.01). (A) APB analysis (B) Quantitative analysis of relative levels of Q-tRNA^His^_GUG_

**Figure 3.**
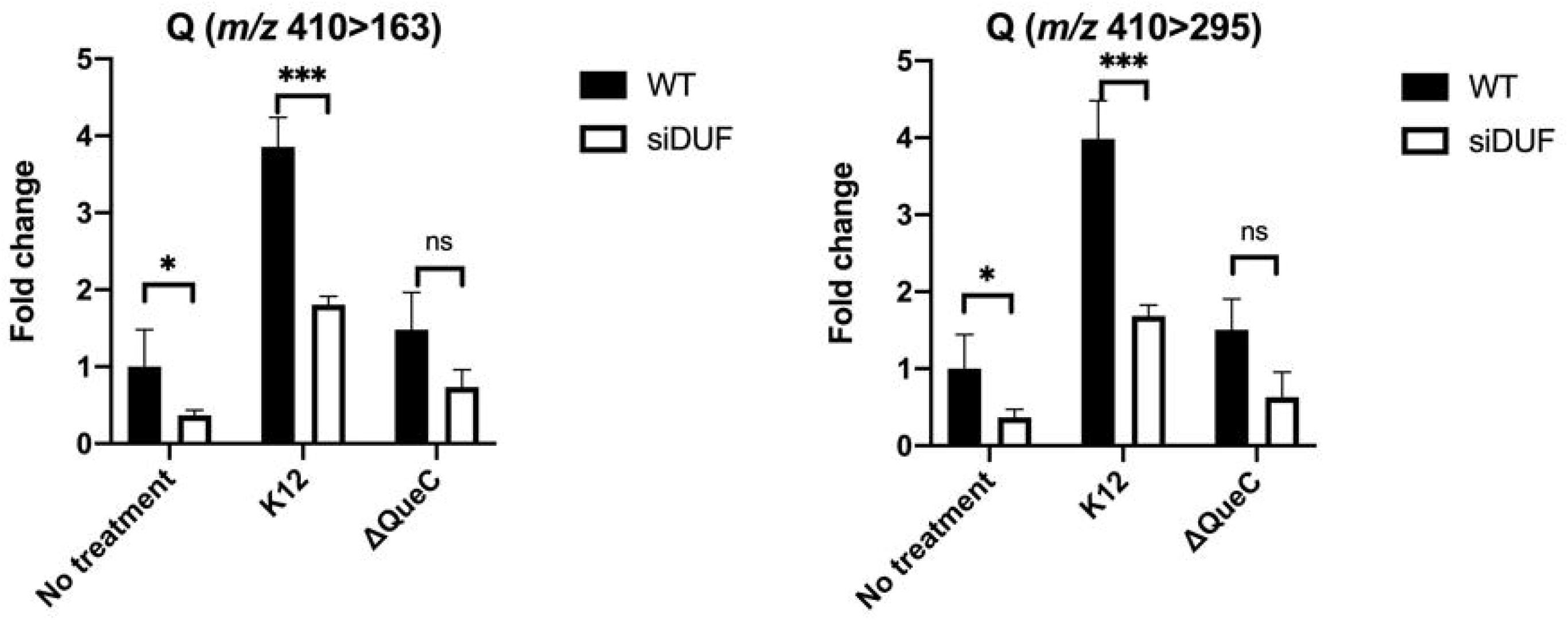
Q level change upon trophozoites cultured with *E.coli* K12 vs. *E.coli* ΔQueC mutants. The fold change is relative to wild type strain without any additive treatment. * indicates *P* value < 0.05, ****indicates *P* value < 0.0001.

### Characterization of EhDUF2419 as a Q-salvaging enzyme in *E. histolytica*

The ability of *E. histolytica* to salvage queuine from Q or from *E. coli* suggests the presence of an active queuine-salvaging pathway in the parasite. We hypothesize that DUF2419 is involved in this pathway. According to annotations of *E. histolytica*’s genome, there is a homologue of *S. pombe* DUF2419 (accession number Q9HDZ9) in *E. histolytica*, namely EhDUF2419 (EHI_098190/XP_653631.1). The EhDUF2419 gene is highly homologous to *S. pombe* DUF2419 (query cover 97%; E value 1E-28; percentage identity 27.1%).

As a first step in the biochemical characterization of EhDUF2419, EhDUF2419 was expressed as a recombinant as GST-tagged protein in *E. coli*. We achieved a level of expression of 100 μg of GST-EhDUF2419 per 100 ml of *E. coli* culture. SDS-PAGE analysis followed by silver staining shows a single 62 kDa band which correspond to the expected molecular weight for GST-EhDUF2419 (Fig 5A). MS analysis confirmed that the 62 kDa band was GST-EhDUF2419 (Fig 5B). The next step was to test the ability of EhDUF2419 to catalyze the formation of queuine from Q. GST-EhDUF2419 was incubated overnight with Q at 37°C and the formation of queuine was determined by LC-MS. When GST-EhDUF2419 was incubated in the presence of Q, significant levels of queuine were detected (Fig 5C). In contrast, no queuine was detected when Q was incubated with GST (Fig 5C). According to these data, EhDUF2419 catalyzes the formation of queuine from Q. The conversion of Q to queuine is not completed. It is possible that the reactions conditions used here are not optimal or EhDUF2419 expressed in *E.coli* is not fully functional. It is also possible that Q isn’t the preferential substrate but rather 5’-QMP as previously suggested by Gunduz and Katze [11].

In order to confirm the role of EhDUF2419 as a queuine-salvaging enzyme, we silenced its expression using antisense small RNAs [29], a method previously used to silence the expression of EhQTRT1 [10]. In this method, a gene-coding region to which large numbers of antisense small RNAs map is used as a ‘trigger’ to silence the gene fused to it. Silencing of EhDUF2419 expression was confirmed by qRT-PCR (Fig 6) of EhDUF2419-silenced trophozoites. Next, we tested the ability of EhDUF2419-silenced trophozoites to salvage queuine from Q or from *E. coli* K12. LC-MS indicate that the level of Q-tRNA is strongly reduced in siEhDUF2419 trophozoites that were grown with Q or with *E. coli* K12. However, siEhDUF2419 trophozoites cultivated in presence of queuine were still able to form Q-tRNAs (Fig 1-4). Interestingly, the level of Q-tRNA in siEhDUF2419 trophozoites that were grown with queuine was lower than the level of Q-tRNA in control trophozoites cultivated with queuine. This result suggests that EhDUF2419 and EhTGT are connected (Fig 1). An examination of Q-tRNA^His^GUG levels by APB polyacrylamide gel analysis also supports these conclusions drawn from LC-MS data (Fig 2).

**Figure 4.**
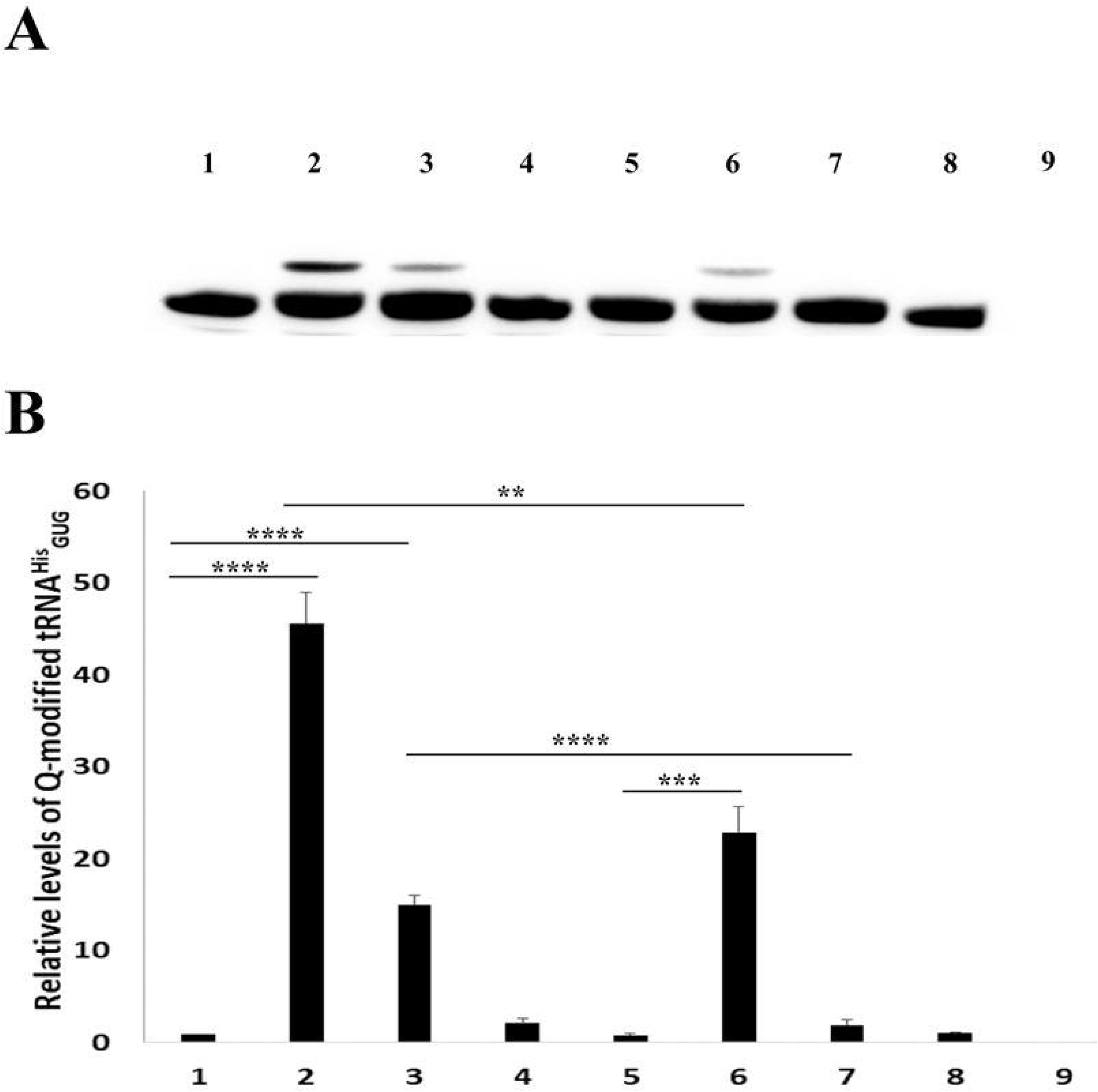
APB northern blot analysis of tRNA^His^_GUG_ in control (WT) and siEhDUF2419 trophozoites that were co-cultivated with *E. coli* K12 or *E. coli* ΔQueC. Control (WT) and siEhDUF2419 trophozoites were cultivated in the presence of *E. coli* K12 or *E. coli* ΔQueC for 7 days (ration of 1 trophozoite: 1000 bacteria). (1) Wild-Type trophozoites (2) queuine-treated WT trophozoites (3) Wild-type trophozoites that were cultivated with *E. coli* K12 (4) Wild-type trophozoites that were cultivated with *E. coli* ΔQueC (5) siEhDUF2419 trophozoites (6) queuine-treated siEhDUF2419 trophozoites (7) siEhDUF2419 trophozoites that were cultivated with *E. coli* K12 (8) siEhDUF2419 trophozoites that were cultivated with *E. coli* ΔQueC (9) *E. coli* K12 RNA. The data represent two independent experiment that were repeated twice. p value<0.05 by an unpaired Student *t* test (**≤0.01, ***≤0.001, ****≤0.0001). (A) APB analysis (B) Quantitative analysis of relative levels of Q-tRNA^His^_GUG_

**Figure 5.**
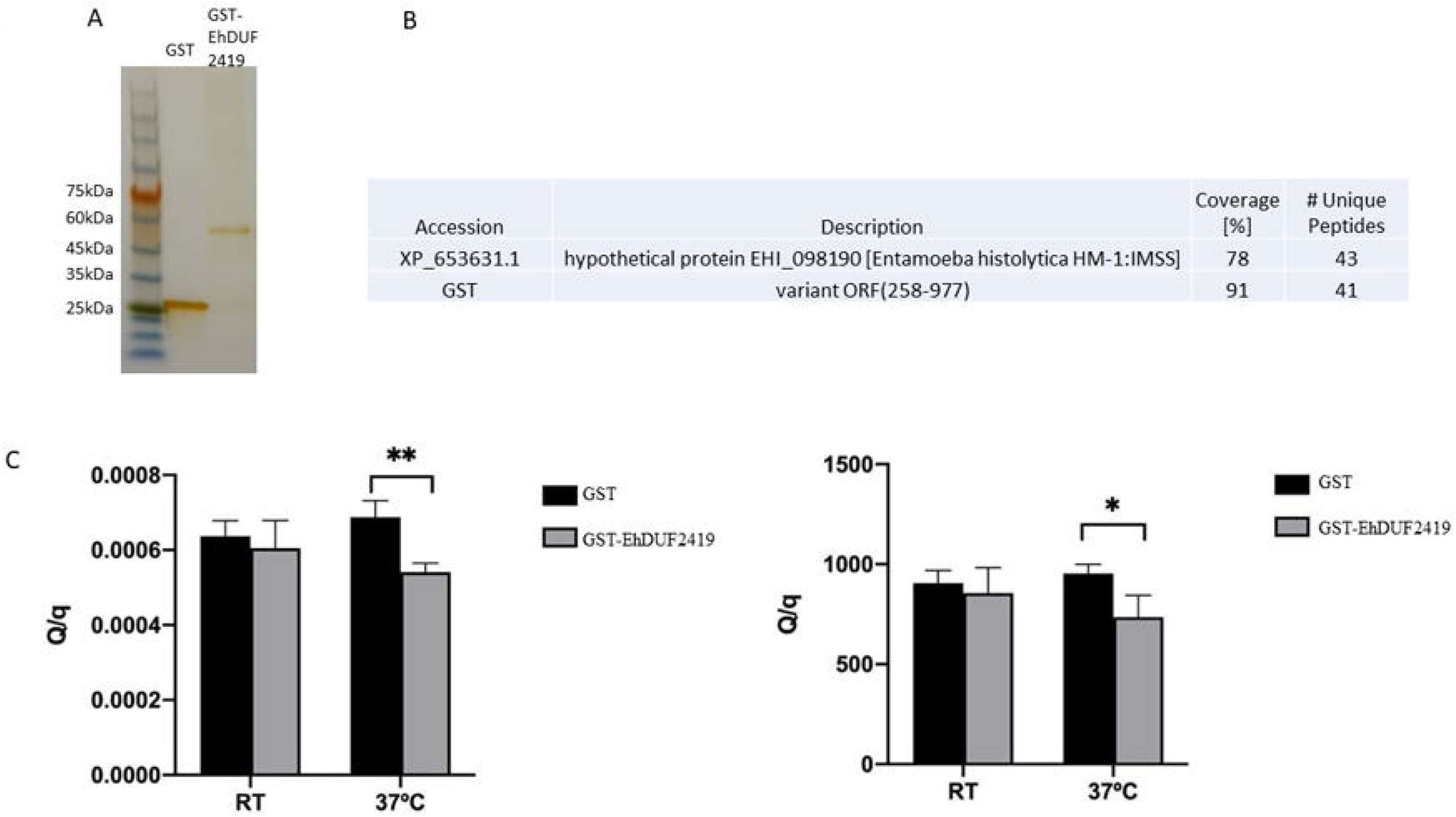
Biochemical characterization of EhDUF2419. **A.** 10 μg of recombinant GST and GST-tagged EhDUF2419 proteins were resolved on 12% SDS gel and stain with silver staining using Pierce Silver Stain Kit according to the manufacturer’s instructions. **B.** Confirmation by MS analysis of the nature of the GST-EhDUF2419 protein. **C.** *In vitro* activity of DUF2419 in Q hydrolysis into queuine. Left panel: ratio of Q UV signal and queuine MS signal; right panel. Ratio of Q MS signal (m/z 410 > 295) and queuine MS signal. * indicated P value < 0.05; ** indicated P value < 0.01.

**Figure 6.**
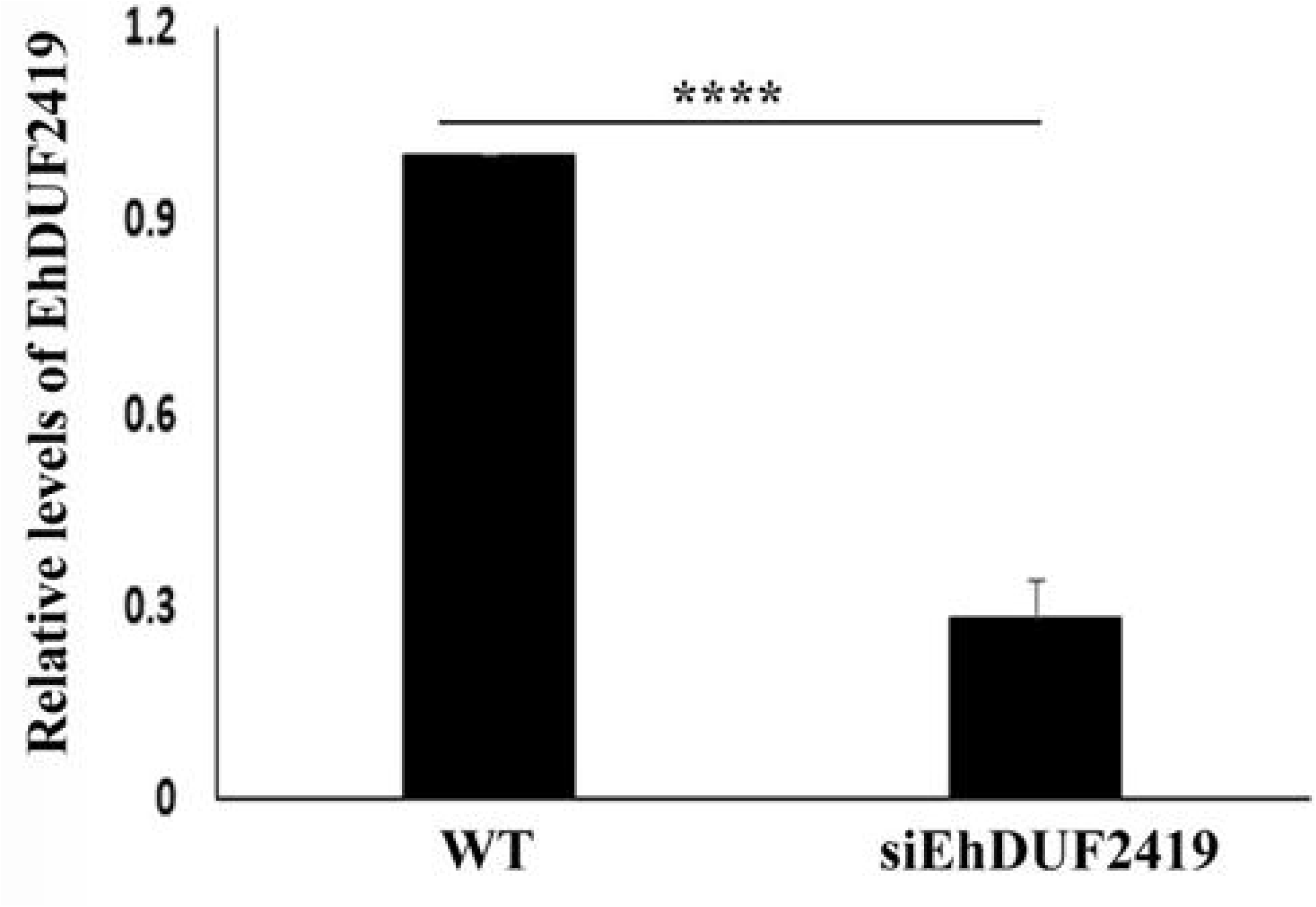
EhDUD2419 expression levels in *E. histolytica* trophozoites. The relative fold change of EhDUD2419 expression in control (WT) and siEhDUF2419 trophozoites was calculated using the 2^^(−ΔΔCt)^ method [19]. The data represent two independent experiments that were repeated twice. p value <0.0001 by an unpaired Student *t* test (****≤0.0001).

As a first step to understand how EhDUF2419 and EhTGT are connected, the levels of EhTGT expression in control and siEhDUF2419 trophozoites was determined by WB analysis. We observed that the EhTGT level in siEhDUF2419 trophozoites is 30% less than in control trophozoites (Fig 7). In this study, we observed that when the expression of EhDUF2419 is downregulated, the expression of EhTGT is also down. It is possible that EhDUF2419 acts as a transcription factor and regulated EhTGT expression. However, except for an homology with DNA glycosidases, no other function can be deduced from DUF2419 sequence [13]. EhDUF2419 may be needed for EhTGT activity but we have already demonstrated that EhTGT is catalytically active without any additional proteins added to the reaction [10]. Finally, EhDUF2419 can also regulates the stability of EhTGT in the parasite. This hypothesis is currently under investigation. Although the link between EhDUF2419 and EhTGT needs more investigation to be understood, the reduction of EhTGT level in siEhDUF2419 trophozoites can explain why less Q-tRNA^His^_GUG_ was observed in siEhDUF2419 trophozoites cultivated in the presence of queuine. We previously reported that the level of Q-tRNAs in the parasite correlates with OS resistance [10]. As less Q-tRNA^His^_GUG_ is formed in siEhDUF2419 trophozoites cultivated in the presence of queuine, the resistance to OS is therefore lower than in control trophozoites cultivated in presence of queuine.

**Figure 7.**
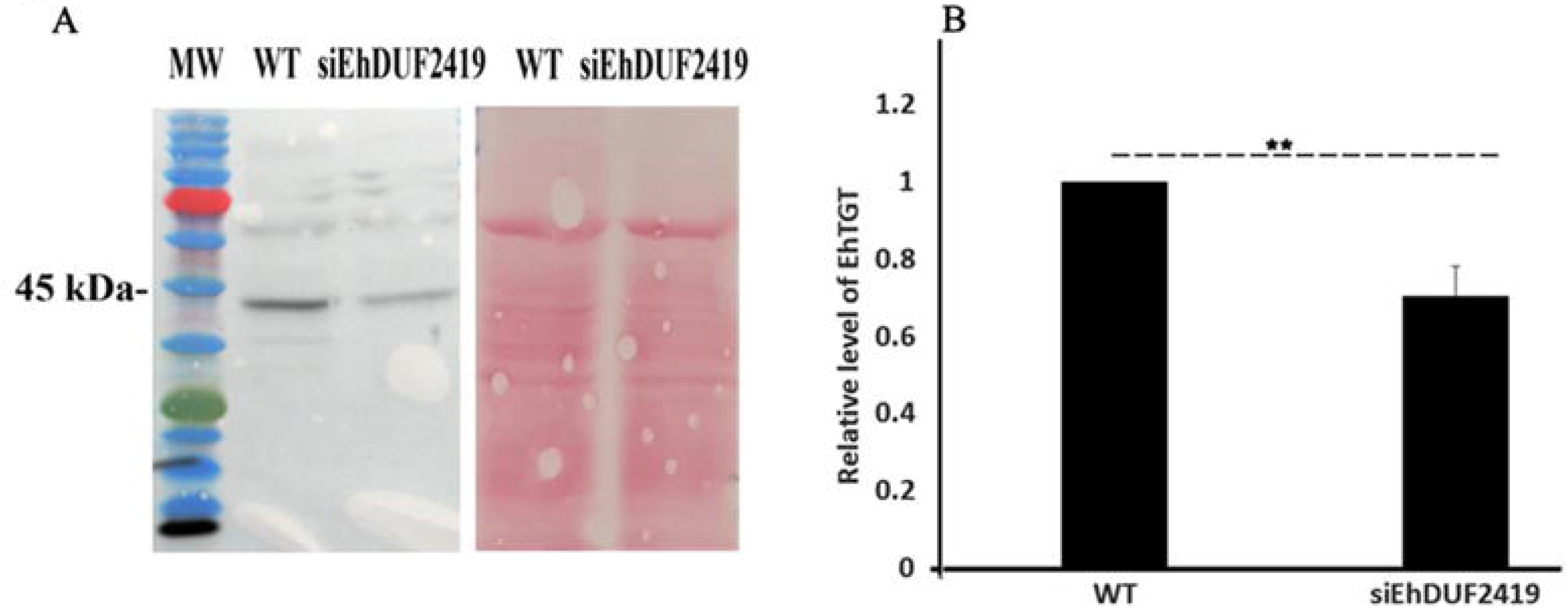
EhTGT level in control (WT) and siEhDUF2419 trophozoites. A. Western blotting was performed on total protein extracts that were prepared from wildtype *E. histolytica* trophozoites (WT) and siEhDUF2419 trophozoites. The proteins were separated on 12% SDS-PAGE gels and analyzed by Western blotting using a homemade EhTGT antibody (1:1,000) [10]. B. Ponceau staining of the membrane before its incubation with EhTGT antibody. The level of EhTGT was normalized according to the total protein amount in each lane as seen by ponceau staining. The experiment was repeated twice. p value <0.05 by an unpaired Student *t* test (**≤0.01).

### Phenotypical characterization of siEhDUF2419 trophozoites

In this study, we examined the effect of silencing EHDUF2419 expression on parasite growth. Our results indicate that this had no effect (supplemental data 1). Our previous work has shown that queuine protects the parasite against OS triggered by H2O2. Here, we have investigated the response of siEhDUF2419 trophozoites exposed to queuine or Q. We observed that queuine but not Q protects siEhDUF2419 trophozoites against OS (Fig 8). In contrast, the level of OS protection achieved by queuine in siEhDUF2419 trophozoites is lower than that observed in control trophozoites.

**Figure 8.**
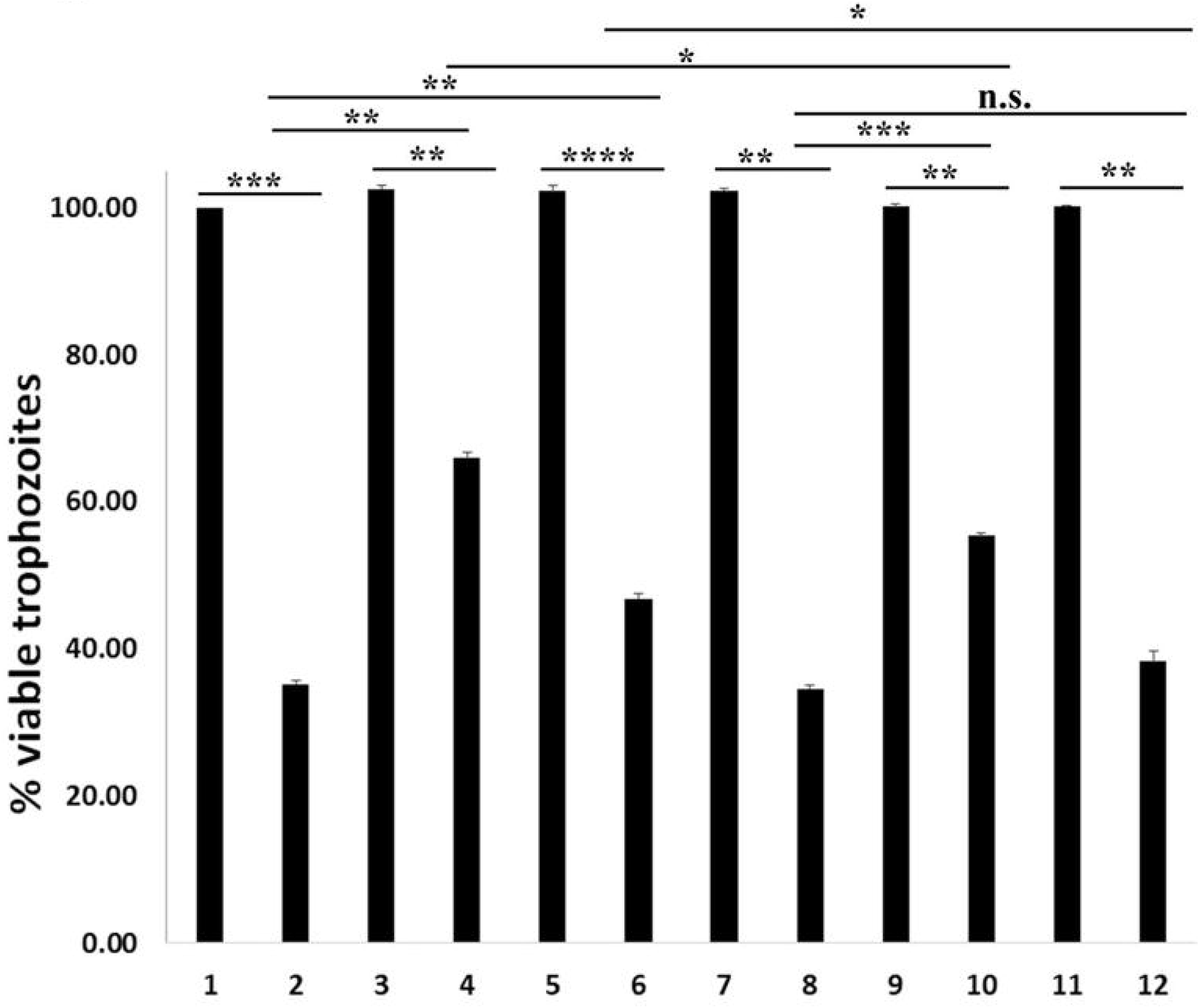
Resistance to OS in queuine/Q treated *E. histolytica* trophozoites. 1x 10^6^ control/siEhDUF2419 trophozoites that grow in the presence of 0.1μM queuine or Q were exposed to 2.5mM H_2_O_2_ for 30 minutes. (1) Control trophozoites (2) Control trophozoites + OS (3) queuine-treated control trophozoites (4) queuine-treated control trophozoites + OS (5) Q-treated control trophozoites (6) Q-treated control trophozoites + OS (7) siEhDUF2419 trophozoites (8) siEhDUF2419 trophozoites + OS (9) queuine-treated siEhDUF2419 trophozoites (10) queuine-treated siEhDUF2419 trophozoites + OS (11) Q-treated siEhDUF2419 trophozoites (12) Q-treated siEhDUF2419 trophozoites + OS. The data represent two independent experiments that were repeated in triplicates. p value<0.05 by an unpaired Student *t* test (*≤0.05, **≤0.01, ***≤0.001, ****≤0.0001).

## Conclusion

In organisms that presumably obtain Q and queuosine from degraded bacteria tRNA material, DUF2419 has been shown to serve as a queuosine salvage enzyme [13]. In this study, we show that DUF2419 is essential for *E. histolytica* to salvage queuosine from phagocytosed bacteria. Obtaining queuosine at the source may help the parasite compete more efficiently for this nutrient with gut bacteria that have been shown to salvage queuosine [4]. We have recently reviewed a number of bacterial metabolites that influence the biology of the parasite including the resistance to OS [30]. Q and queuine which are protecting *E. histolytica* against OS (this work and [10]) represent additional bacterial metabolites that may help the parasite to survive in the large intestine when dysanerobiosis occurs as a result of inflammatory conditions [31] or from dysbiosis [32]. By targeting the salvage pathway identified in this study, a new strategy may be found to control the development of this parasite in the host

## Acknowledgments

This work was supported by the Israel Science Foundation (3208/19), the Ministry of Science and Technology, Israel (1020546) and Niedersachsen-Deutsche Technion (ZN 3454).

