## Supplementary figures and images for "Queuine salvaging in the human parasite *Entamoeba histolytica*"

### Supplemental Fig 1

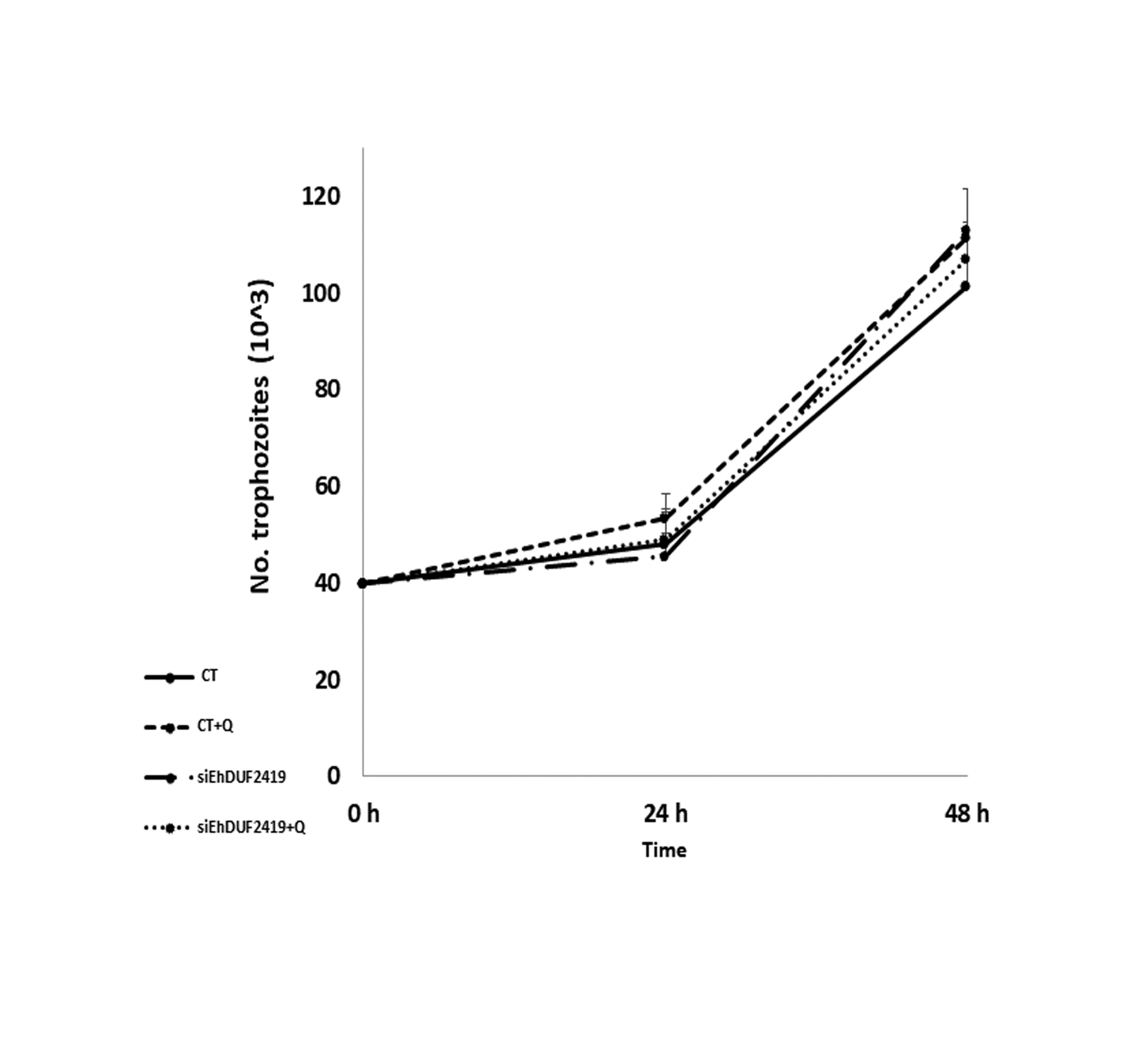
